# Emergent hydrodynamic synchronization between microbeads labeling bacterial flagellar motors

**DOI:** 10.1101/2025.10.19.683311

**Authors:** Tsubasa Ishihara, Nariya Uchida, Shuichi Nakamura

## Abstract

Synchronization is a fundamental phenomenon observed across a wide range of physical, chemical, and biological systems. Even under the low-Reynolds-number conditions that govern microbial motility, synchronization of rhythmic processes is considered essential for diverse activities ranging from single cells to populations, yet experimental evidence remains scarce. Here, we performed rotation measurements of microbeads attached to two truncated bacterial flagella and observed the emergence of phase synchronization between these nanoscale active motors. The resulting intermittent in-phase synchronization was analyzed using a hydrodynamic model incorporating elastic deformation of the flagella, showing that stronger hydrodynamic coupling promotes more stable phase-locking. This study experimentally demonstrates fluid-mediated biological synchronization using bacterial rotary machinery, deepening our understanding of the physical principles underlying rhythmic phenomena and self-organization in living systems.

## Introduction

Synchronized motion of periodic locomotor organs, such as legs, wings, and flagella, underlies directional locomotion across biological systems. In animals, such coordination is typically achieved through the interplay of sensory information and neural circuits [1, 2], while vortex-mediated interactions also induce phase matching in bird flocks [3] and fish schools [4–7].

In contrast, microorganisms lack a nervous system, and their synchronization must arise from fundamental physical principles. Indeed, at microscopic scales―where viscous forces dominate and inertia is negligible synchronization is known to emerge from purely hydrodynamic interactions. A classical example is the metachronal wave generated by coordinated beating of cilia on the surface of *Paramecium*, which drives forward swimming through a traveling wave of net flow [8]. In the green alga *Chlamydomonas*, two eukaryotic flagella beat synchronously in a breast-stroke-like pattern [9, 10]. Such synchronous beating of cilia and flagella has long been attributed to hydrodynamic coupling, a mechanism first proposed by Taylor in his seminal theoretical work [11]. Recent studies, however, suggest that synchronization may also involve non-hydrodynamic factors. For example, the metachronal coordination in *Paramecium* is influenced by cell surface elasticity [12], and body rocking contributes to flagellar synchronization in *Chlamydomonas* [13].

Bacterial flagella, by contrast, are fundamentally different from their eukaryotic counterparts. Each flagellum consists of a helical filament that functions as a screw-like propeller, rotated by a membrane-embedded flagellar motor (Figure 1a). The flagellar motor is driven by the ion motive force, combining membrane potential and trans-membrane ion gradients ― protons in *Escherichia coli* and *Salmonella enterica*, or sodium ions in *Vibrio* species [14]. The flagellar motor is composed of a rotor and a dozen stators, and the translocation of coupling ions through the ion channels formed within the individual stators is converted into torque to rotate the rotor. The torque generated at the basal body is transmitted to the helical filament via the short, flexible joint called the hook. While some bacteria, such as *Vibrio cholerae* and *Pseudomonas aeruginosa*, possess a single polar flagellum, peritrichous bacteria like *E. coli* and *S. enterica* carry multiple flagella per cell. Peritrichous bacteria swim by bundling flagella into a cohesive structure for smooth, forward motion (run) and reorient by unbundling them (tumble). The run-tumble pattern underlies chemotaxis ― allowing bacteria to respond to gradients of chemicals, temperature, or oxygen ― and is essential for their survival strategy [14].

**Figure 1.**
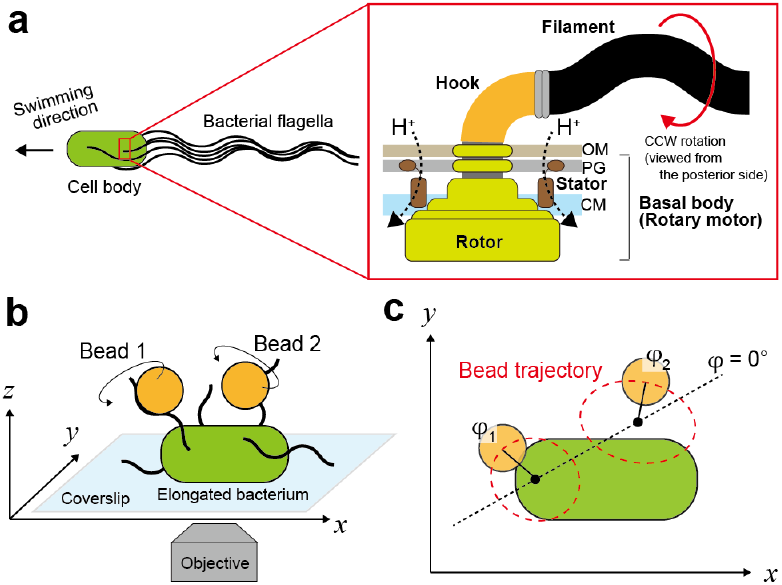
Schematic diagrams of the bacterial flagellum and the experimental setup. (a) A peritrichous bacterium (left) and a crosssectional illustration of the flagellar basal body (right). The motor is embedded in the cell envelope, consisting of the outer membrane (OM), peptidoglycan layer (PG), and cytoplasmic membrane (CM). Energy from proton translocation through membrane-bound stators is converted into torque. The motor is connected to a left-handed helical filament via a flexible hook that functions as a universal joint. While the motor is bi-directional, counterclockwise (CCW) rotation generates the thrust required for forward swimming. (b) The dualbead assay configuration used to investigate the synchronization of two adjacent flagella. (c) Definition of *ϕ*_1_ and *ϕ*_2_ in the bead assay.

Although synchronization between bacterial flagella has not been observed, its significance has been discussed theoretically, particularly in the context of flagellar bundling. During bacterial swimming maneuvers, multiple flagella form a coordinated bundle, as depicted in Figure 1a. To explain how these independent motors achieve the necessary coordination, theoretical studies have explored various aspects, such as the effects of torque [15], the number of flagella [16], inter-flagella distance [17], and flagellar flexibility [18, 19] on the synchronization and bundling of flagella. However, despite decades of theoretical predictions, direct experimental evidence of such synchronization in peritrichous bacteria remains remarkably scarce.

The most established method for analyzing bacterial flagellar dynamics is the bead assay. This technique has elucidated fundamental characteristics of the flagellar motor at the single-unit level, including its torque-speed relationship, stator assembly dynamics, and mechanochemical coupling mechanisms [20, 21]. Previous studies using multiple beads have assessed coordinated behaviors, such as the sequential switching of motor direction associated with the diffusion of chemotaxis-signaling molecules [22] or speed changes occurring nearly simultaneously in two motors in response to the propagation of energy variation [23]. In contrast, the problem of instantaneous phase synchronization via extracellular hydrodynamic coupling has not been addressed.

Here, we performed rotation measurements of microbeads attached to two truncated bacterial flagella to investigate the emergence of phase synchronization between these nanoscale rotary motors in a living cell (Figure 1b, c). Operating in a high-load regime ― where the torque is less sensitive to speed fluctuations and the stator units remain stably incorporated ― achieves a robust physical experimental system. This setup revealed intermittent inphase synchronization between the beads, with the stability of the phase-locking sensitive to the inter-bead distance and the frequency mismatch between the motors. A theoretical model, incorporating the elastic deformation of the flagella and hydrodynamic coupling between the rotating beads, quantitatively accounted for the observed behaviors. These findings provide direct evidence for hydrodynamically mediated synchronization in the bacterial flagellar motor, bridging the unprecedented dynamics of living molecular motors with the framework of nonlinear physics.

## Results

### Flagellar synchronous rotation

We carried out a bead assay to measure rotations of two *Salmonella* flagella simultaneously (Figures 1b and 2a). We determined the phases of two beads attached to truncated flagellar filaments (*ϕ*_1_ and *ϕ*_2_) from their rotation trajectories (Figure 1c) and then determined the temporal variation of phase difference between two beads (Δ = *ϕ*_1_ − *ϕ*_2_). The time courses of bead positions and Δ (Figure 2b) show intermittent in-phase rotations between the two beads, interrupted by occasional phase slips. The kymograph analysis also represents phase coincidence and phase slip between two beads (Figure 2c) .

**Figure 2.**
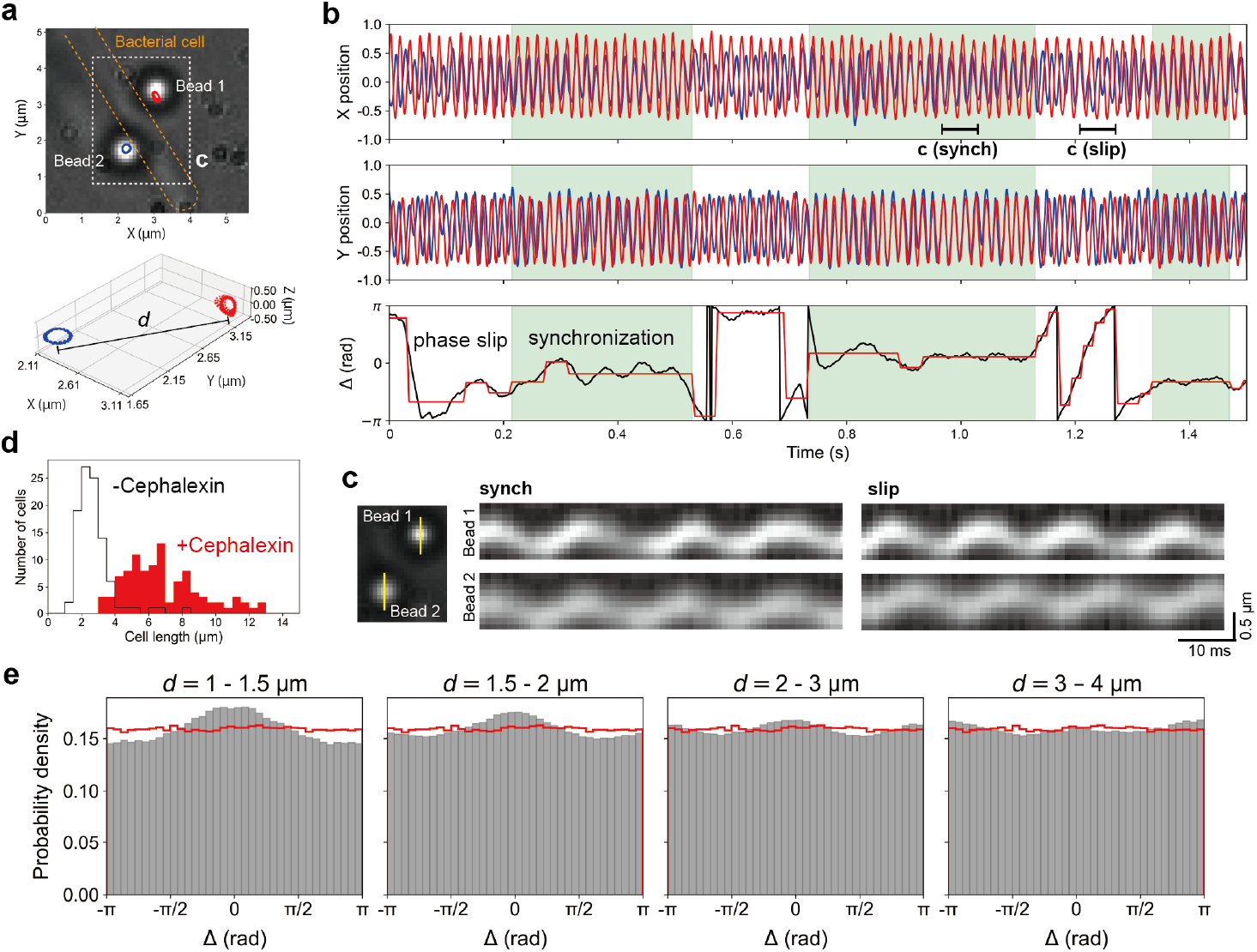
Simultaneous rotation measurements of two neighboring beads. (a) 1-*µ*m beads attached to flagella extending from the same bacterial cell (upper) and 3D rotation trajectories (lower). The addition of cephalexin elongated the bacterial cells. Distances between beads (*d*) were defined as those between rotational centers. The area indicated by a white box is analyzed in (c). (b) Time variations of bead positions (X and Y) are shown in the upper panel. Data from two beads are discriminated by blue and red. The phase differences (Δ) determined for the time variations of bead positions are shown in the lower panel. The results of stepwise smoothing (see Figure S4c) are shown in red. The periods of in-phase synchronization are highlighted in green. See Figures S1 and S2 for additional example data. The kymographs of the periods designated by horizontal bars in the X position are shown in (c). See also Movie 1. (c) Kymographs along the yellow lines in the left panel show in-phase synchronization (synch) and phase slip (slip) between Bead 1 and Bead 2. (d) Effect of cephalexin on the cell length. (e) Dependence of the phase-difference distribution on *d*: 10, 46, 28, and 13 pairs were analyzed for *d* =1-1.5 *µ*m, 1.5-2 *µ*m, 2-3 *µ*m, and 3-4 *µ*m, respectively. The red lines represent the results for”pseudo pairs” created by randomly pairing beads that were either located sufficiently far apart or observed in different chambers. To assess accidental synchronization, beads with approximately equal intrinsic frequencies were selected for the pseudo pairs.

The addition of cepharexin to the growth media elongated the bacterial body length (Figure 2d), enabling us to investigate if the distance between beads (*d*) affects their rotations. Analysis of the phase difference distributions obtained from the rotation traces revealed a prominent peak near Δ = 0, indicating in-phase synchronization, and the peak height decreased with an increment of *d* (Figure 2e). We note here that if the two beads rotate at similar speeds, such in-phase synchronization could arise by chance, even in the absence of any interaction. To control for this possibility, we generated “pseudo pairs” by randomly combining rotation traces of beads that were either spatially well-separated on different cells or obtained from independent experiments conducted in separate observation chambers (see Figure S3 for example rotation data). The peak-distribution analysis of pseudo pairs showed no peak at any specific phase difference for these pseudo pairs (red lines in Figure 2e). These results suggest that neighboring beads experience distance-dependent interactions that intermittently induce in-phase synchronization.

### Variation of the intrinsic-frequency difference

We extracted the periods during which the two beads were asynchronized and determined their respective intrinsic frequencies, *ω*_1_ and *ω*_2_ (Figures S4a, b). Focusing on the difference between *ω*_1_ and *ω*_2_ (= Δ*ω*), we detected multiple peaks in the distribution that were approximately equally spaced (Figure 3a). ∼ 10 stator units are incorporated into the flagellar motor, and they are known to turn over during rotation [24]. Though the stator assembly is expected to be relatively stable in the present experiment performed at high load, the number of stators can vary between bacterial cells and motors and may change over time. Such variations of the stator numbers have been observed as a motor-speed distribution with multiple peaks and stepwise changes in motor speed [25, 26]. Rotation assays of the *Salmonella* flagellar motor have shown speed distributions and stepwise speed variation with an interval of 6 - 8 Hz [26], which is consistent with the peak intervals observed in the Δ*ω* (red arrows in Figure 3a). There may have been a difference of up to four stator counts between the motors analyzed as a pair. The Δ*ω* due to this difference in stator number also had a distinct effect on the synchronization of the two beads, showing that the larger the Δ*ω*, the smaller the synchronous time fraction (Figure 3b). Although the variation in stator number may have been accentuated by the extension of the cell length with cephalexin, synchronization occurs when the stator number differs by only one or two.

**Figure 3.**
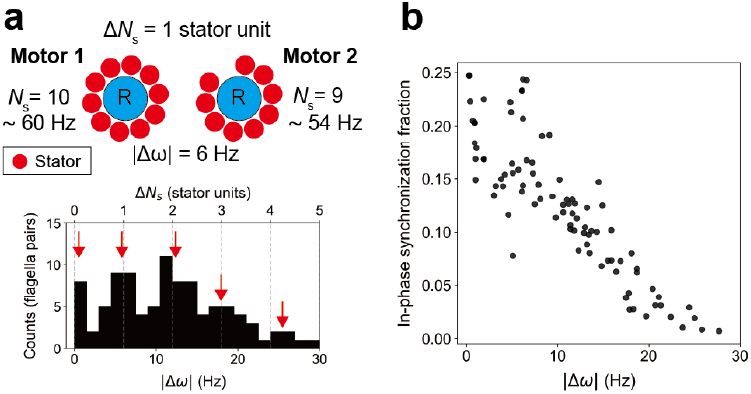
Difference in intrinsic frequencies. (a) Relationship between the number of stators and flagellar motor speed. The flagellar rotor (R) is surrounded by approximately 10 stators. Discrete differences in flagellar rotation rates between motor pairs are associated with differences in the number of stator units (see *main text*). For example, if the speed difference (|Δ*ω*|) between Motor 1 and Motor 2 is 6 Hz, the stator-number gap (Δ*N*_s_) is estimated to be 1 unit (upper panel). The lower panel shows the distribution of |Δ*ω*| (*n* = 85 motor pairs). Red arrows indicate approximate peak positions. The Δ*N*_s_ values estimated from |Δ*ω*| are indicated on the upper axis, and vertical dashed lines are drawn at 6 Hz intervals. (b) Dependence of the in-phase synchronization fraction on |Δ*ω*|. The in-phase synchronous fraction is the ratio of the sum of in-phase synchronous periods, determined from Δ vs time plots as shown in Figure 2b, to total measuring time. Each dot represents a pair of beads.

### Synchronization model introducing elastic deformation

Our dual-bead assay revealed that neighboring flagella rotate in phase, often sustained for several tens of revolutions. What mechanism underlies the synchronization of this bacterial nanomachine? To address this ques-tion, we focused on hydrodynamic interactions between oscillators ― a possibility originally proposed by Taylor in the context of low Reynolds number environments [11]―and conducted a theoretical investigation. In the case of *Chlamydomonas*, breaststroke-like swimming is achieved through a repeated cycle of an effective stroke, where the flagella push fluid forcefully, and a recovery stroke, which slowly returns them to the initial position. A representative class of theoretical models assumes that such periodic modulations of driving torque can lead to hydrodynamic synchronization [27] (see also [28, 29] for reviews). However, although bacterial flagellar torque exhibits stochastic fluctuations, there is no clear evidence for periodic modulation. Another model treats active fluid flow along the flagellar axis as the source of synchronization in bacterial carpets, where the cells are attached to the substrate [30]. However, in the present setup, flow induced by the rotating tethered cell is more important than the flagella-generated axisymmetric flow. This is similar to the situation of elastic cilia, where distortion of the trajectories modulates hydrodynamic interaction and induces synchronization [31]. Building on the previous work, we developed a new theoretical model that emphasizes the role of flagellar elasticity in enabling synchronization.

In our model, each tethered cell is represented by a bead that is driven by a constant force and makes a circular trajectory in the absence of hydrodynamic flow (Figure 4). The flagellar elasticity is mimicked by a harmonic spring that anchors each bead to the center of the circle. The spring is extended by the hydrodynamic forces and adjusts its length on a timescale much shorter than the rotation period. Therefore, the positions of the beads are well described by their phases *ϕ*_1_, *ϕ*_2_ only. The flow field generated by the rotating beads is described by the Stokes law for the drag force and the Blake tensor, the Green function of the Stokes equation in the presence of a no-slip boundary. In studying synchronization, we focus on the time evolution of the phase difference Δ = *ϕ*_1_ − *ϕ*_2_. Incorporating small differences in the natural rotation frequencies *ω*_1_, *ω*_2_ and the gyration radii *R*_10_, *R*_20_ of the two beads, we obtain 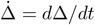 in the form

**Figure 4.**
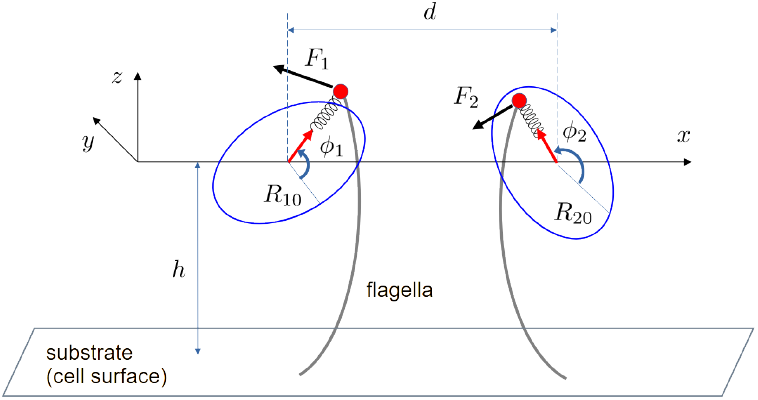
Hydrodynamic model. The beads represent the tethered cells. Their positions are specified by the phases *ϕ*_1_, *ϕ*_2_ and the radial displacements from the circular trajectories of radii *R*_10_ and *R*_20_. See the text and Supplementary Information for details.

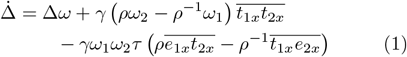

Here, *γ* = 9*ah*^2^*/d*^3^ is a geometrical coupling parameter, where *a* is the bead radius (with *a* = 0.5 *µ*m) and *h* is the height of the rotational orbit from the surface (∼ 1 *µ*m). *ρ* = *R*_20_*/R*_10_ is the ratio of the intrinsic orbital radii, *τ* is the radial relaxation time, *t*_*ix*_ and *e*_*ix*_ are the *x*-components of the radial and tangential unit vector of the intrinsic orbits, and *Ā* means as cycle average of any quantity *A*. For the circular orbits, we obtain

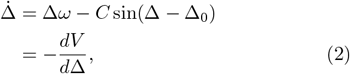

where *C* (*>* 0) and Δ_0_ are constants determined the parameters mentioned above, and

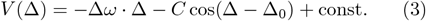

is an effective potential [27]; see Supplementary Information for details of the model. Note that Eq.(2) has a stationary (i.e., phase-locking) solution only if |Δ*ω*| ≤ *C*.

### Phase-locking stability

We assessed the variation rate of Δ, 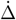, from Δ vs time plots. 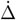 was calculated approxi-mately every one revolution: 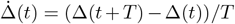, where the period *T* was defined as 20 ms based on an estimated rotation frequency of approximately 50 Hz. The obtained distributions of 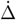 show convex downward with a minimum around Δ = 0 (Figure 5a and c). 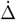 can be described using the effective potential *V* (Δ) as 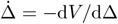, allowing us to estimate the potential landscape for the flagellar synchronization experimentally. As predicted from the 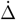 distribution, *V* (Δ) shows the potential minimum at Δ = Δ_0_ ∼ 0 (Figure 5b and d), suggesting the stability of the in-phase synchronous state.

**Figure 5.**
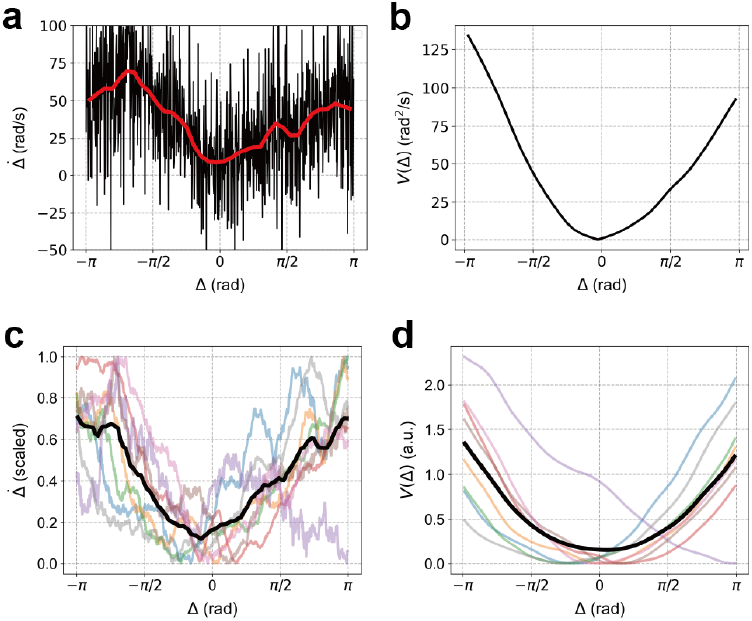
Effective potential inferred from time-varying phase differences. (a) Distribution of 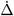 (see *main text*) against Δ obtained from rotation data of a pair of beads. The red line represents the moving average of the black plots (median of the raw data). (b) An effective potential estimated from (a). (c) 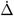 vs Δ plots of multiple different pairs of beads scaled by individual minimum values (light colored lines) and their average plot (thick black line). (d) An effective potential estimated from (c) and their average plot (thick black line).

According to calculations that account for the elasticity of the bacterial flagellar filament (∼ 3.5 pN · *µ*m^2^) [32] and the hook in the torsionally loaded state (∼ 3.0 pN · *µ*m^2^) [33], together with their lengths and the surrounding viscosity, the time constant *τ* of flagellar synchronization is estimated to be ≈ 0.3 ms (Supplementary Information). Although the filament-specific, elasticity-dependent fluctuations in the rotational trajectory that the model predicts to precede synchronization were not clearly resolved, intermittent disturbances of the rotational path were frequently observed during rotation (Figure 2a and b). Measurements with higher spatiotemporal resolution are expected to clarify the relationship between these trajectory disturbances and synchronization.

### Synchronization by hydrodynamic coupling

In our model, the hydrodynamic force acts only along the direction connecting the centers of the two orbits (the *X*-axis in Figure 4), and its magnitude is described by the third term in the time-evolution equation of Δ (hereafter referred to as the interaction term). The position vectors of the beads, the ratio of their orbital radii, and the frequency difference included in this interaction term were experimentally measured in this study. The geometrical factor (*γ*) and the radial relaxation time of the beads (*τ*) can be estimated from the bead-assay setup and from literature values (Supplementary Information, see [32–34] for physical properties of flagella). By averaging this experimentally determined interaction term over one period (assuming Δ remains constant within a cycle), we obtain *C* sin Δ (Eq.(2)). In other words, when the interaction term is plotted against sin Δ as in Figure 6a, the slope provides the coefficient *C*, which represents the interaction strength (hereafter referred to as the coupling coefficient). Since the interaction term has the dimension of inverse time (s^−1^), *C* also has the dimension of s^−1^. Thus, multiplying *C* by the oscillation period *T* (determined from the mean intrinsic frequency of each bead pair) yields the dimensionless coupling coefficient *CT* . We then examined the relationship between *CT* and the time fraction of inphase synchronization observed for each bead pair, and found that larger values of *CT* correlated with longer synchronization times (Figure 6b).

**Figure 6.**
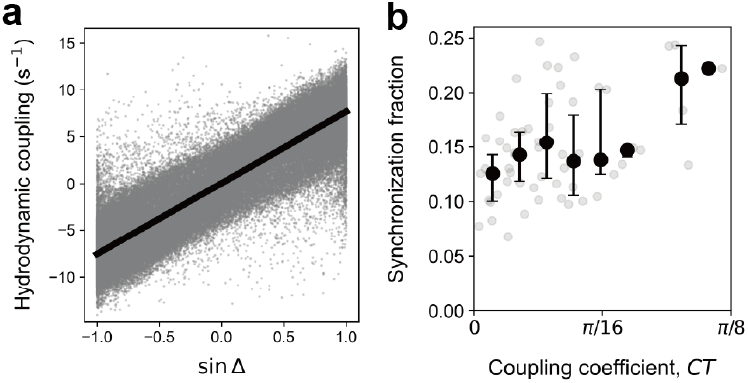
Hydrodynamic coupling. (a) Relationship between hydrodynamic coupling strength and phase difference. The gray dots are values obtained from a time-series data of a pair of beads, and the black line is the result of line fitting. (b) Dependence of in-phase synchronization on the coupling coefficient. The dimensionless coupling coefficient *CT* is plotted (see *main text* for details). The gray dots indicate data from individual bead pairs. The black dots and error bars represent the median and interquartile range within each bin, where the range 0 to *π/*8 was divided into ten equal bins.

According to the time-evolution equation of Δ, phase locking (i.e.,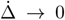) requires |Δ*ω*| ≤ *C* as mentioned in the model section. Experimentally, bead pairs with larger |Δ*ω*| were indeed less likely to synchronize (Figure 3b). For example, a typical pair with intrinsic frequencies near 60 Hz and Δ*ω* = 6 Hz (roughly one stator difference) would require a coupling strength of *CT >* 0.2*π* to synchronize, since the phase slip per period is Δ*ω* · *T* = 6 Hz · (1*/*60 s) · 2*π* = 0.2*π*. Our analysis included pairs with |Δ*ω*| up to 15 Hz (corresponding to a difference of less than three stators), which would theoretically require *CT >* (15*/*6) · 0.2*π* = *π/*2 for every pair to be synchronized. Therefore, our estimation *CT < π/*8 is consistent with the observed partial synchronization with the time fraction *<* 1*/*4 (i.e., ∼0.25 at its maximum in Figure 6b).

Furthermore, the model assumes that radial relaxation is much faster than the rotation period, requiring *ωτ* ≪ 2*π*. Given the flagellar elasticity and viscous drag coefficient, we estimate *τ* ∼ 0.3 ms (Supplementary Information). For *ω* ∼ 60 Hz, this gives *ωτ* ∼ 0.02, which satisfies this requirement under our experimental conditions. Taken together, these findings suggest that the hydrodynamic interaction assumed in the theory is indeed the emergent factor underlying synchronization of bacterial flagella.

Theoretically, the coupling constant *C* scales with the bead distance *d* and height from the cell surface *h* as *C* ∝ *h*^2^*/d*^3^ (via *γ*), while its dependence on the orbital radius *b* is negligible in our approximation (see the Supplementary Information). This assumption is justified by our experimental data, which show a narrower distribution of *b* than *d* and small *b/d* ratios (Figure S6a). Considering the relatively constant *b* and *h*, we tested the scaling law *CT* ∝ 1*/d*^*n*^. This yielded a best fit with *n* = 2.82 (Figure S6b), which is in excellent agreement with the theoretical prediction *n* = 3. This result strongly supports the validity of the long-distance approximation in our model. On the other hand, a large scatter in *CT* was observed at short distances (Figure S6b), which might reflect nontrivial dependencies on the specific orientations of the two orbits [35] that are not captured by our simplified model. Elucidating these effects is a subject for future work and will require experiments with higher spatiotemporal resolution.

### Flow around rotating beads

Having established that hydrodynamic interactions can quantitatively account for the observed synchronization, we next sought to directly visualize the underlying flow field. To this end, we introduced tracer beads into the chamber during observations of beads rotating on bacterial flagella and tracked their trajectories. Tracking tracer beads around a single rotating bead revealed a counterclockwise flow pattern consistent with the direction of rotation (Figure 7a). When tracer beads were introduced during observations of two beads rotating in close proximity on flagella protruding from the same bacterium, a continuous counterclockwise streamline encompassing both beads was visualized (Figure 7b). These observations suggest that the continuous flow connecting the two beads could contribute to the hydrodynamic coupling underlying their phase-locked rotation.

**Figure 7.**
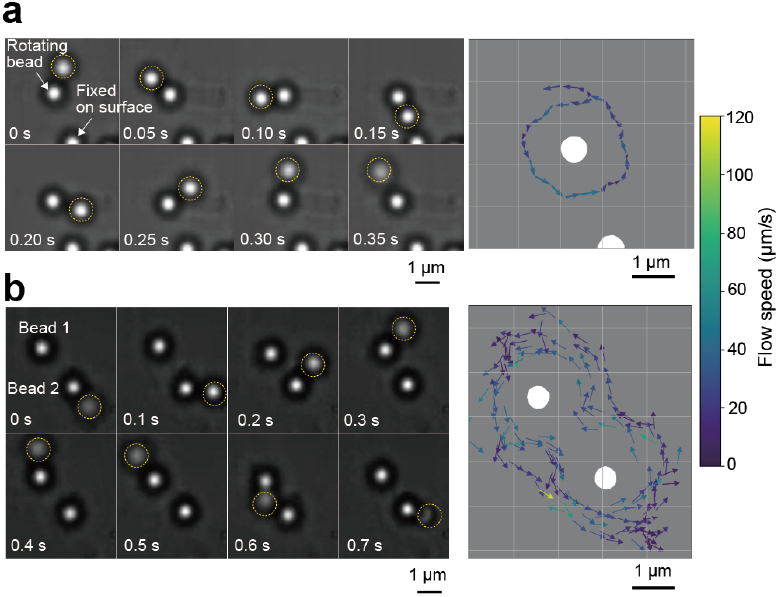
Flow around rotating beads. (a) Movement of a free bead (circled by a yellow dashed line) around a single rotating bead attached to a flagellum (left) and flow estimated from the free-bead movement (right). (b) Movement of a free bead around two rotating beads and the estimated flow. See Movie 2 and Movie 3 for a single bead and paired beads, respectively.

## Discussion

Our rotation measurements of two flagellar motors using bead probes revealed intermittent in-phase synchronization when the motors rotated in close proximity. The emergence of synchronization was found to be sensitive to both the inter-bead distance and the intrinsic frequency mismatch between the two motors ― the latter likely arising from variations in the number of stators assembled into each motor. Our hydrodynamic model, which incorporates changes in the rotational trajectory induced by flagellar filament elasticity, confirmed that inter-bead fluid interactions drive the propensity for synchronization. To our knowledge, these findings represent the first experimental demonstration of synchronization in a rotary molecular motor operating within a living cell.

Bacterial flagella are undoubtedly capable of in-phase synchronization, but does such intermittent coordination confer any advantage to the bacterium? The most plausible physiological relevance lies in the process of flagellar bundle formation (Figure 1a). Preceding numerous innovative theoretical studies, macroscale model experiments and simulations have suggested that in-phase synchronization is essential for enabling stable bundling [36, 37].The bending rigidity of bacterial flagella has been reported to lie in the range of 2–4 pN · *µ*m^2^, corresponding to a Young’s modulus of up to ∼ 1 GPa [38, 39]. This modulus is comparable to cytoskeletal proteins in biological systems and to soft plastics or Teflon in engineered materials [40]. Such intrinsic stiffness suggests that constraints on the relative phase between rotating filaments are likely critical for maintaining the structural integrity of the bundle. Recent theoretical work has proposed that hydrodynamic interactions within the flow fields generated by bacterial motion can promote flagellar bundling [41], suggesting that directed locomotion contributes substantially to this process. In this context, transient episodes of inphase rotation might play a critical role in nucleating or stabilizing the initial stages of bundling, while allowing for subsequent adjustments in motor torque or filament orientation.

However, one must remain cautious when extrapolating phenomena observed in truncated flagella labeled with 1 *µ*m beads to probe-free, full-length flagellar filaments. Regarding the load on the motor, the non-linearity of the torque-speed relationship must be considered: in the highload regime, the torque remains relatively insensitive to speed fluctuations, whereas in the low-load regime, the torque decreases sharply as speed increases [20]. While 1 *µ*m beads operate in the high-load regime, probe-free flagellar filaments belong to the low-load regime. This difference in output characteristics may significantly influence the motor’s response to hydrodynamic interactions. Furthermore, the assembly stability of the stators is sensitive to load; lower loads facilitate stator dissociation [42–44]. In our experiments, relatively strong synchronization was induced when the difference in stator numbers was within 1–2. In contrast, for flagellar filaments in the low-load regime, frequent stator turnover might hinder stable synchronization. Conversely, compared to the interactions between spherical beads, flagellar filaments are expected to exhibit longer-range and stronger axial hydrodynamic coupling. Thus, linking the present physical results to physiological significance requires a careful consideration of load regimes and probe scales.

In summary, we have successfully applied a well-established biophysical assay to demonstrate the phase synchronization of active flagellar motors in living cells and analyzed the resulting dynamics within a quantitative dynamical systems framework. Whereas locomotion at high Reynolds numbers relies heavily on neural control, fluid-mediated synchrony at low Reynolds numbers serves as a fundamental physical mechanism ensuring the robust operation of microorganisms’ machinery. The synchronization of bacterial flagellar motors identified here represents a novel instance of coupled active oscillators, further deepening our understanding of the physics underlying rhythmic phenomena in living systems.

## Methods

### Bacteria and media

*S. enterica* serovar Typhimurium strain MM3076iC [(Δ*cheA*–*cheZ*), fl*iC* (Δ204–292)] was used, which possesses flagellar motors that exclusively rotate counterclockwise with sticky flagellar filaments to be labeled with latex beads [45]. L-broth (LB) and motility medium were prepared as described previously [46]

### Bead assay

Bacteria grown overnight in LB at 37^◦^C were diluted 1:100 into fresh LB and incubated for 3h at 37^◦^C with shaking. After adding 20 *µ*g/mL cefalexin to elongate the cell length [47], the mixture was further incubated for 1h at 37^◦^C with shaking. Bacteria collected by centrifugation were suspended into a motility medium and their sticky flagellar filaments were truncated by passing through a 27-gauge needle. The bacterial suspension was infused into a glass chamber to adhere the bacteria to a glass surface, and 1-*µ*m latex beads were infused into the chamber. Unbound beads floating in the chamber were removed by infusing motility medium, and bead rotation was observed under a bright-field microscope (BX53, 100 × oil immersion objective, Olympus). Rotations of two beads attached to two filaments of the same cells were recorded by a high-speed CMOS camera (IDP-Express R2000, Photron) at intervals of 1 ms and their angular positions were determined using the line connecting the two rotation centers as the origin. See Supplementary Information for details of data analysis.

## Data Availability

The data supporting the findings of this study are available from the corresponding author upon request.

## Code Availability

The computer codes used for this study are available from the corresponding author upon reasonable request.

## Acknowledgements

We thank Dr. S. Toyabe (Tohoku University) for providing software for rotation analysis. This work was supported by the JSPS KAKENHI: 22H04828 and 24K02274 for SN, and 24K0985 for NU.

## Author contribution

SN and NU designed the project. TI and SN set up the optical system and performed the experiments. NU performed theoretical research. All authors wrote the manuscript and approved the submitted version.

