## Supplementary Information and Figures for "Emergent hydrodynamic synchronization between microbeads labeling bacterial flagellar motors"

Title: Direct observation of emergent hydrodynamic synchronization between bacterial flagella

1. Example paired beads rotation measurement (Figure S1, Figure S2)
2. Example single-bead rotation measurement (Figure S3)
3. Determination of intrinsic frequencies (Figure S4)
4. Reconstruction of 3D coordinates from 2D trajectories (Figure S5)
5. Derivation of the model
6. Parameter estimation
7. Validity of the long-distance approximation in experiments (Figure S6)

SI Video 1. A pair of beads attached to flagella extending from the same bacterium intermittently exhibits in-phase synchronization (Synch) and phase slips (Slip).

SI Video 2. The flow around a rotating bead attached to a bacterial flagellum was visualized by tracking the movement of a freely floating bead in the chamber.

SI Video 3. The flow around the rotating paired beads attached to flagella extending from the same bacterium was visualized by tracking the movement of a freely floating bead in the chamber.

### 1 Example paired beads rotation measurement

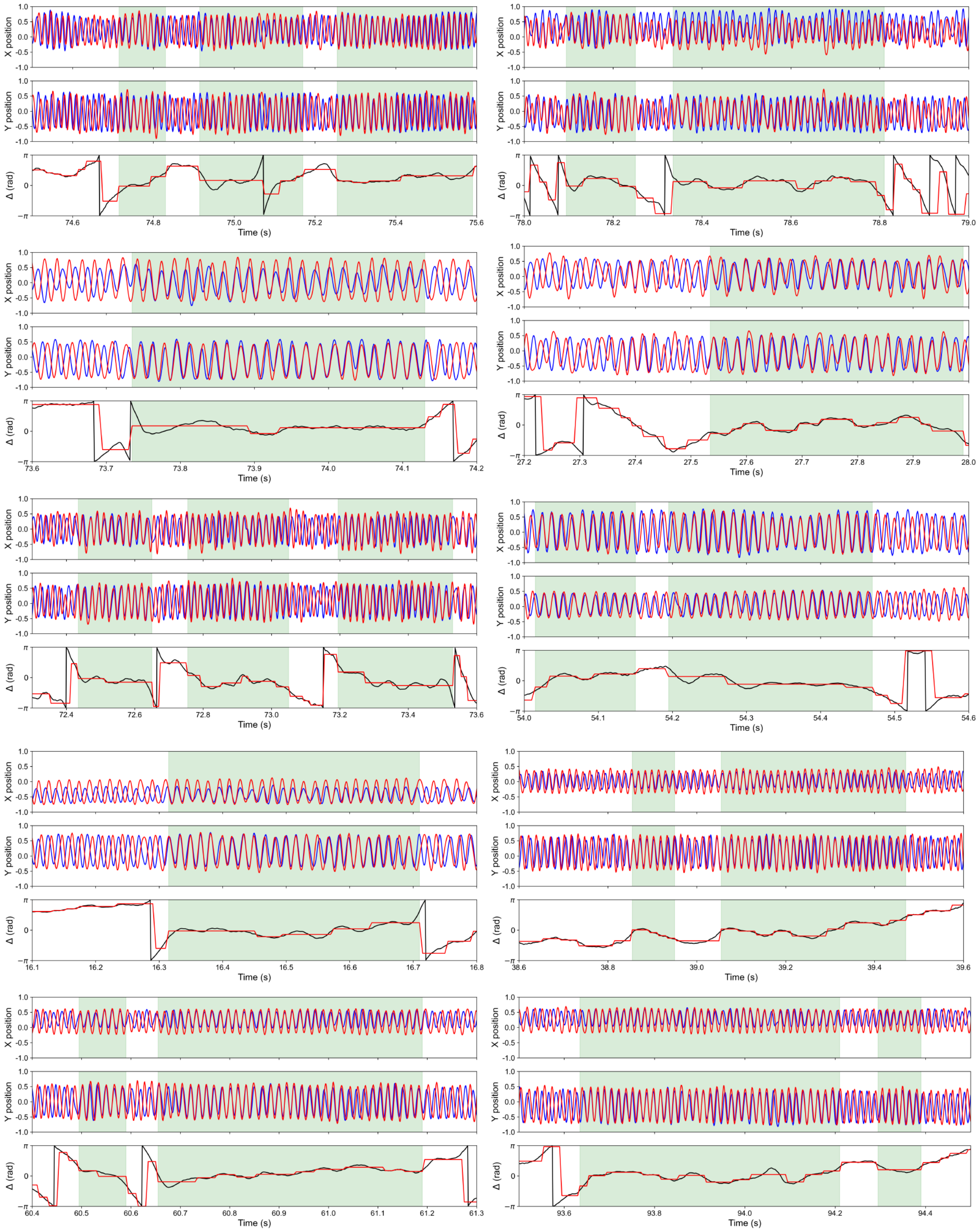

Figure S 1. See Figure S2 for explanation.

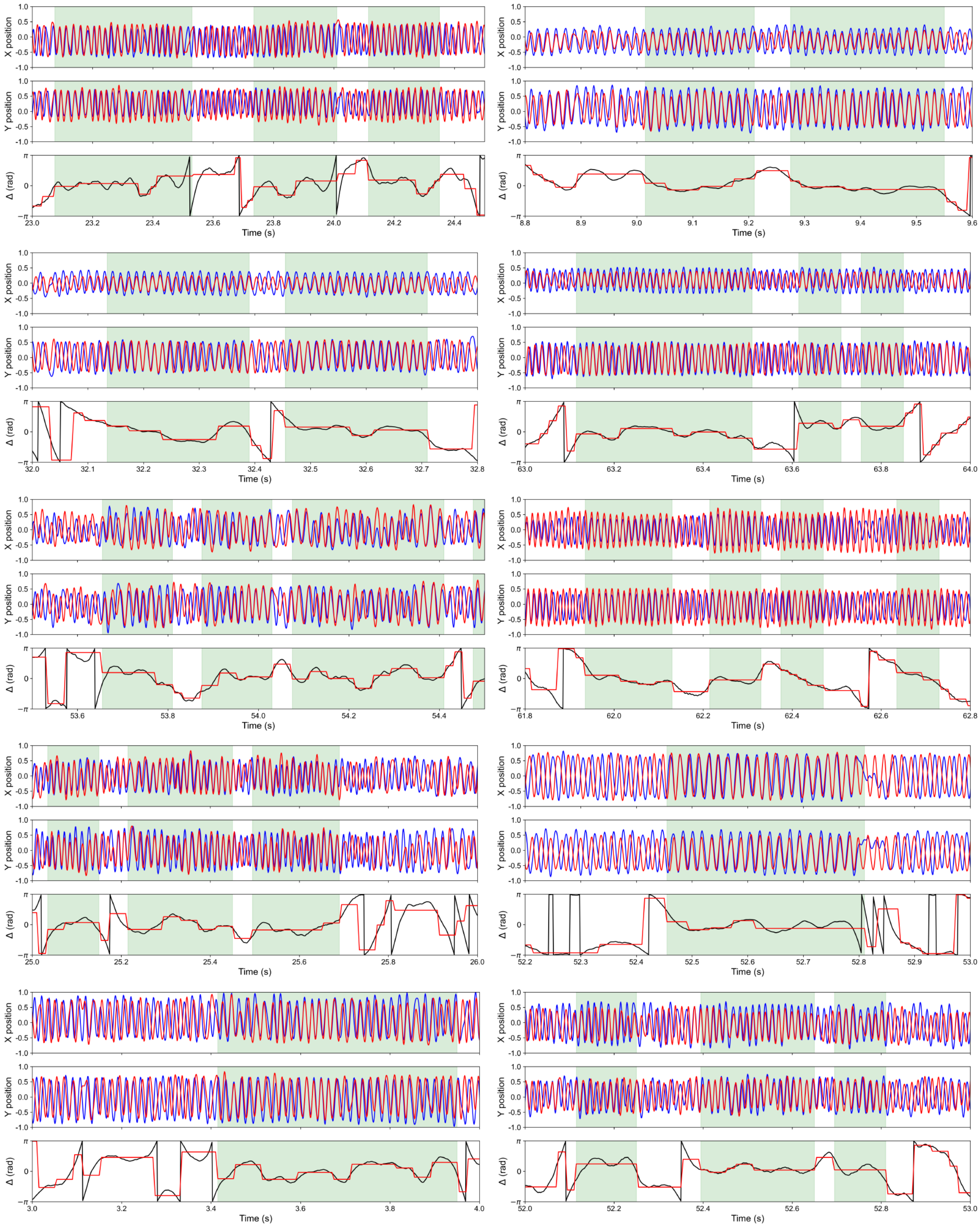


Figure S 2. Continued from Figure S1. We aligned 20 pairs of rotational measurement data, each covering 1 s (10 pairs in Figure S1 and 10 pairs in Figure S2). As in Figure 2 in the *main text*, the red and black traces represent the centroid displacements of the two paired beads (arbitrarily labeled bead1 and bead2). The angular difference calculated from the bead centroids was statistically processed (red line, method described later), and intervals identified as in-phase synchronization are highlighted in green.

### 2 Example single-bead rotation measurement

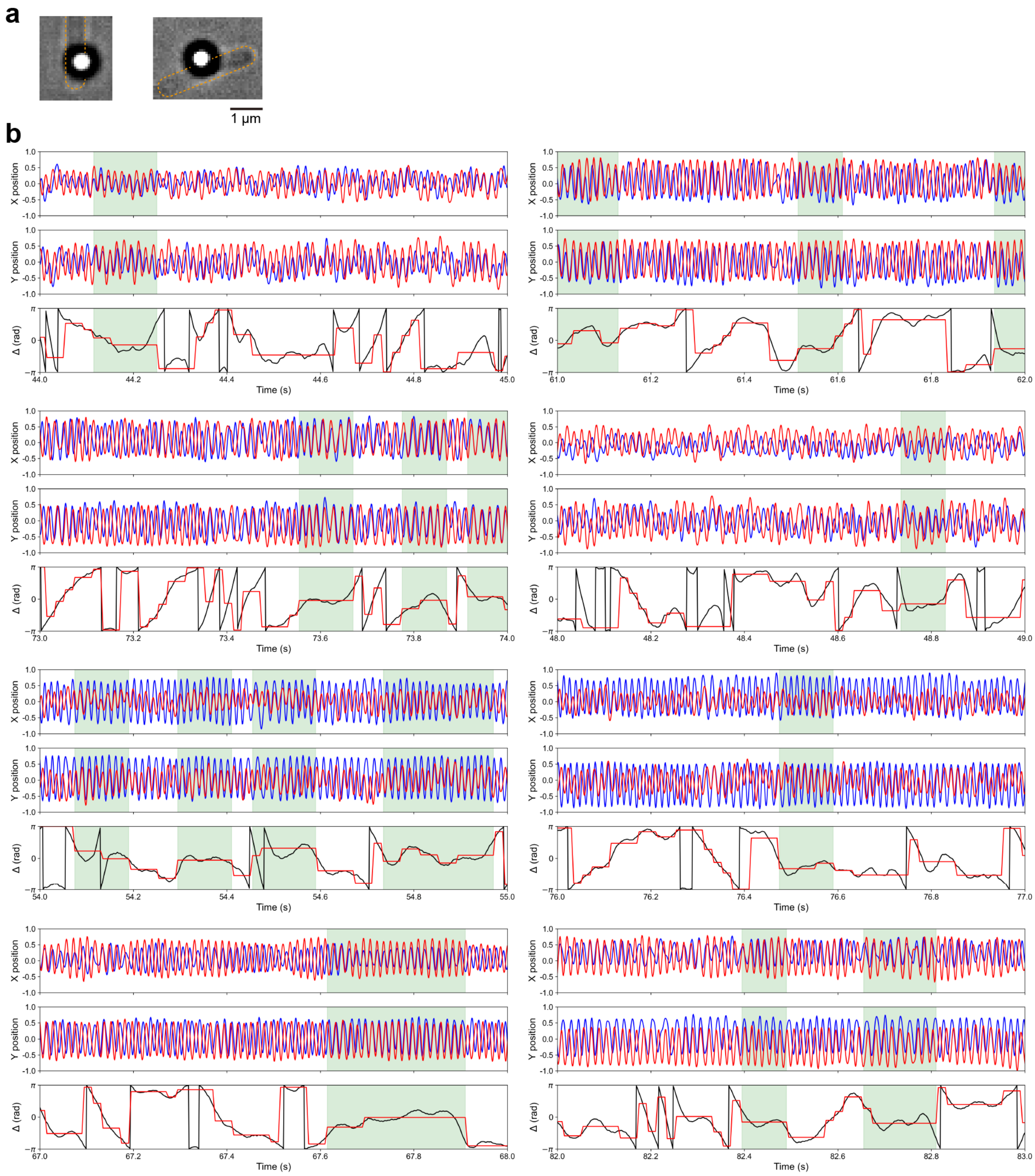


Figure S 3. Example single-bead rotation measurement. (a) Micrographs of a bead attached to a single flagellar filament. (b) Results from examining accidental synchronization by combining beads with close intrinsic frequencies that were measured either sufficiently far apart or in separate chambers. As in Figures S1 and S2, periods where the rotational phases coincide are highlighted in green. Since no interactions exist between the two beads, the phase synchronization observed here is an accidental phenomenon resulting from their close intrinsic frequencies. The intrinsic frequencies of the analyzed pairs were as follows: from top to bottom in the left column, 60 Hz, 63 Hz, 70 Hz, and 63 Hz; and from top to bottom in the right column, 63 Hz, 60 Hz, 70 Hz, and 63 Hz. Because these beads were unaffected by the rotation of other beads during measurement, the average of all velocity values obtained over the entire measurement period (approximately 100 s) was taken as the intrinsic frequency of each bead (motor).

### 3 Determination of intrinsic frequencies

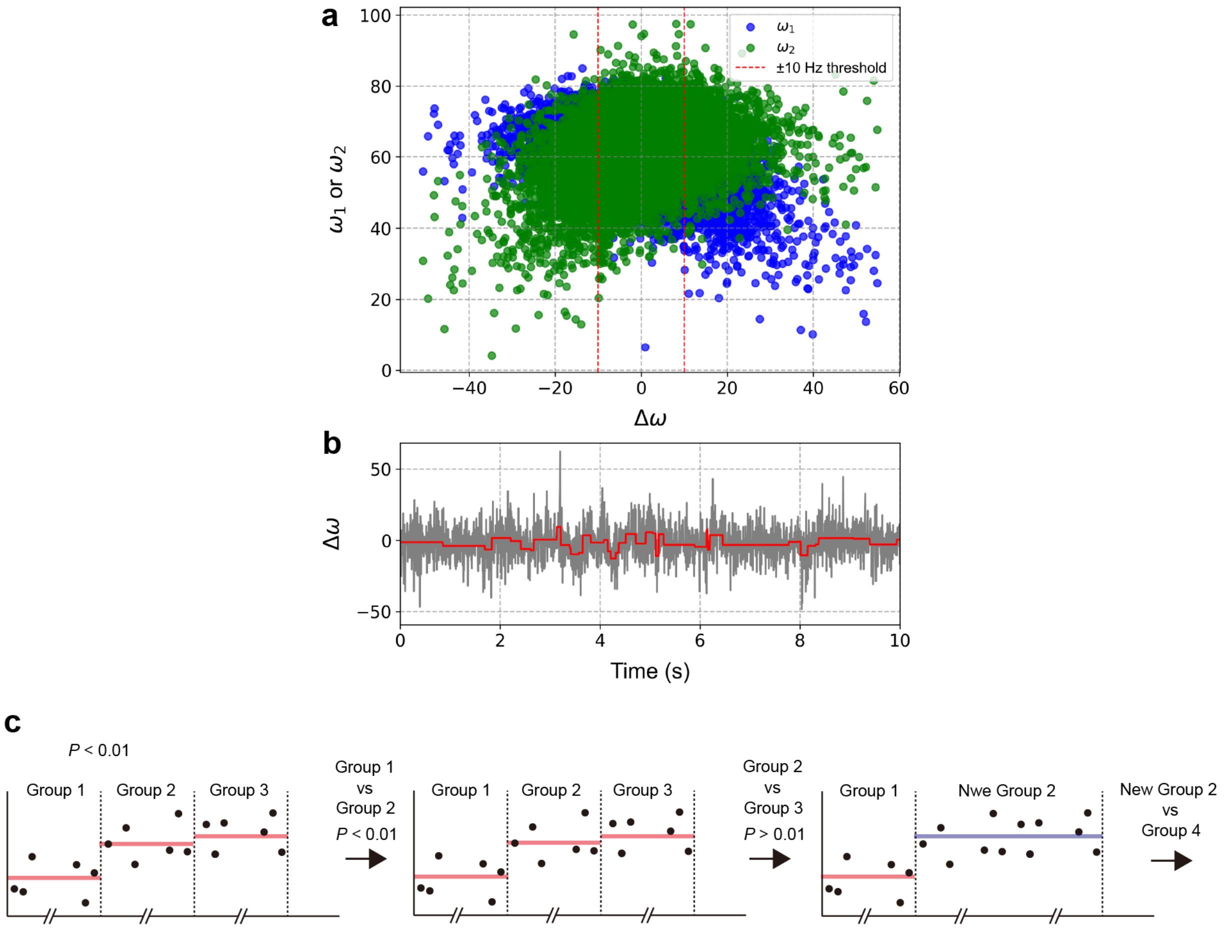


Figure S 4. Determination of intrinsic frequencies in paired beads. (a) Rotation rates of two beads ($\omega_1$ and $\omega_2$), determined at 5-ms intervals, were plotted against the speed difference ($\omega_1 - \omega_2 = \Delta\omega$). Because beads rotating in close proximity are always subject to some degree of fluid-mediated interaction, it is difficult to determine their true intrinsic frequencies. We also attempted an experiment in which one bead was stopped using an optical trap; however, the trap influence extended to the neighboring bead as well, preventing independent and accurate determination of the intrinsic frequency. In this study, we calculated the rotational velocity every 5 ms from the time series of bead centroids measured at 1 ms intervals, and obtained the velocity difference between the two beads at each 5 ms step. Although the rotational speed of the bacterial flagellar motor fluctuates due to various factors, one major source of large changes is the turnover of stator units [1]. For flagella attached to 1-$\mu$m beads, the contribution of a single stator to the motor speed is typically 5–10 Hz [2, 3]. Based on this, periods in which the velocity difference remained below 10 Hz were regarded as synchronized and excluded from intrinsic frequency determination. In other words, the mean velocity during periods where the absolute velocity difference exceeded 10 Hz was defined as the intrinsic frequency. The dashed red lines in *a* represent the threshold of 10 Hz. (b) The time variation of $\Delta\omega$. The raw data and the result of statistical processing are shown in gray and red, respectively. (c) The statistical processing, shown in red in (b). The time-series data, recorded at 1-ms intervals, were first divided into windows of four data points (20 ms each $\sim$ 1 revolution for a bead rotating at 50 Hz). Adjacent windows were then statistically compared: if a significant difference was detected, the windows were kept separate; if not, they were merged into a single window. This comparison and merging procedure was repeated over the entire dataset until all neighboring windows were statistically distinct. This statistical analysis was also used to determine the duration of synchronization and asynchronization.

### 4 Reconstruction of 3D coordinates from 2D trajectories

The raw data consist of the time series of bead centroids projected onto the $XY$ plane. First, the two-dimensional trajectories were smoothed by low-pass filtering and cubic spline interpolation. For each time window, an ellipse was fitted to the projected coordinates using a least-squares method. From the major and minor axes of the ellipse, $a$ and $b$, the eccentricity was used to estimate the tilt of the rotation plane. Specifically, the inclination angle $\phi$ of the plane was calculated as

$$\phi = \arccos\left(\frac{a}{b}\right). \tag{1}$$

The coordinates were then rotated to align the ellipse with the coordinate axes:

$$x' = x\cos\theta + y\sin\theta, \tag{2}$$
$$y' = -x\sin\theta + y\cos\theta, \tag{3}$$

where $\theta$ is the rotation angle obtained from the ellipse fitting.

Finally, the $z$-coordinate was reconstructed by scaling the transformed $x'$ coordinate according to the tilt angle:

$$z = \pm x' \sin\phi, \tag{4}$$

with the sign determined by the orientation of the major axis. The rotation center was fixed at $z = 0$.

In this way, the two-dimensional bead trajectories were converted into three-dimensional coordinates, which were then used for further analysis and visualization.

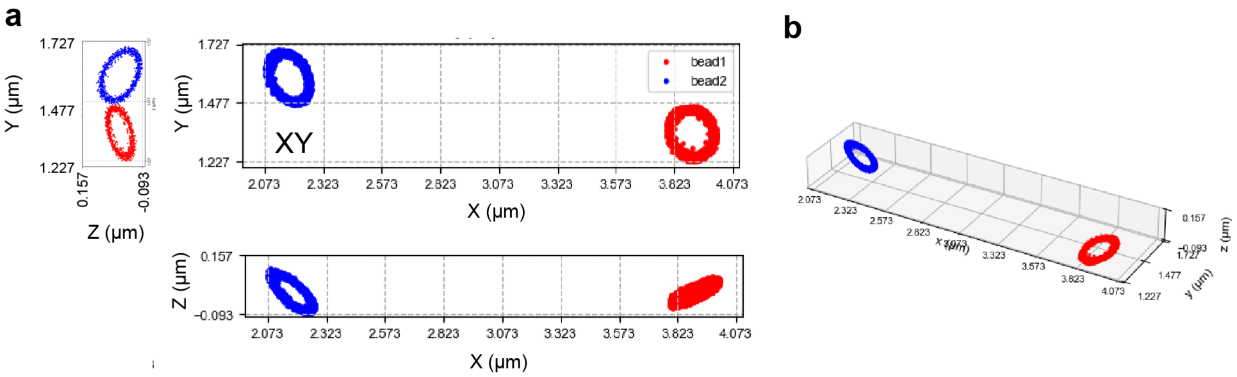


Figure S 5. Reconstruction of 3D coordinates from 2D trajectories.

### 5 Derivation of the model

First we define the two reference (circular) trajectories $C_1$, $C_2$ and the phases $\phi_1$, $\phi_2$ of the two beads. As shown in Figure 4, we assume that the centers of $C_1$ and $C_2$ are located at the same height $h$ from the substrate ($z = -h$) and on the $x$-axis with the distance $d$. The position $\boldsymbol{r}_i$ of the bead on each trajectory is given by

$$\boldsymbol{r}_i = \boldsymbol{r}_{i0} + \boldsymbol{Q}_i \cdot (R_i \cos\phi_i, R_i \sin\phi_i, 0)^{\mathrm{T}}, \qquad (i = 1, 2), \tag{5}$$

where $\boldsymbol{r}_{i0}$ is the center position of $C_i$, $R_i(t)$ is the distance from the center, $\boldsymbol{Q}_i$ is the rotation matrix that maps a circle on the $xy$-plane to the tilted trajectory. The matrix also defines the unit vectors

$$\boldsymbol{n}_i = \boldsymbol{Q}_i \cdot (0, 0, 1), \tag{6}$$
$$\boldsymbol{e}_i = \boldsymbol{Q}_i \cdot (\cos\phi_i, \sin\phi_i, 0), \tag{7}$$
$$\boldsymbol{t}_i = \boldsymbol{Q}_i \cdot (-\sin\phi_i, \cos\phi_i, 0), \tag{8}$$

which give the torque direction, the radial direction and the tangential direction, respectively. The position and velocity of the $i$-th bead is thus expressed as

$$\boldsymbol{r}_i = \boldsymbol{r}_{i0} + R_i \boldsymbol{e}_i, \tag{9}$$
$$\dot{\boldsymbol{r}}_i = \dot{R}_i \boldsymbol{e}_i + R_i \dot{\phi}_i \boldsymbol{t}_i. \tag{10}$$

The orbiting motion of the bead is driven by the tangential driving force $\boldsymbol{f}_i^{(d)} = F_i \boldsymbol{t}_i$, where force magnitude $F_i$ is assumed constant. The flagellar elasticity is represented by a harmonic spring of spring constant $k$ and natural length $R_{i0}$, which anchors the bead to the center of $C_i$. Therefore, the spring exerts the elastic force $\boldsymbol{f}_i^{(e)} = -k(R_i - R_{i0})\boldsymbol{e}_i$ on the bead. The bead exerts the force $\boldsymbol{f}_i^{(v)} = \zeta[\dot{\boldsymbol{r}}_i - \boldsymbol{v}(\boldsymbol{r}_i)]$ on the surrounding fluid, where $\zeta = 6\pi\eta a$ is the Stokes drag coefficient ($\eta$: shear viscosity, $a$: bead radius) and $\boldsymbol{v}(\boldsymbol{r})$ is the velocity field governed by the Stokes equation $\eta\nabla^2 \boldsymbol{v} = \sum_i \boldsymbol{f}_i^{(v)}$ with the no-slip boundary condition ($\boldsymbol{v} = \boldsymbol{0}$) at the substrate. In analogy with electrostatics, the no-slip boundary condition is treated by introducing an image force that constitutes a force dipole when combined with the original force. We obtain

$$\boldsymbol{v}(\boldsymbol{r}_i) = \sum_{j \neq i} \boldsymbol{G}(\boldsymbol{r}_i - \boldsymbol{r}_j) \cdot \boldsymbol{f}_i^{(v)} \tag{11}$$

where $\boldsymbol{G}(\boldsymbol{r})$ is the Green function known as the Blake tensor [4]. Assuming $d \gg h, R_i$, we can render it to the simple form [5]

$$\boldsymbol{G}(\boldsymbol{r}_i - \boldsymbol{r}_j) \approx \frac{\gamma}{\zeta} \boldsymbol{e}_x \boldsymbol{e}_x, \qquad \gamma = \frac{9ah^2}{d^3}, \quad \boldsymbol{e}_x = (0, 0, 1). \tag{12}$$

The dimensionless constant $\gamma (\ll 1)$ measures the strength of the hydrodynamic coupling. Note that the inverse-cubic dependence on the distance is characteristic of dipole-dipole interactions. The force balance equation $\boldsymbol{f}_i^{(d)} + \boldsymbol{f}_i^{(e)} + \boldsymbol{f}_i^{(v)} = 0$ is decomposed into the radial and tangential parts as

$$\zeta \left[\dot{R}_i - \boldsymbol{v}(\boldsymbol{r}_i) \cdot \boldsymbol{e}_i\right] = -k(R_i - R_{i0}), \tag{13}$$
$$\zeta \left[R_i \dot{\phi}_i - \boldsymbol{v}(\boldsymbol{r}_i) \cdot \boldsymbol{t}_i\right] = F_i. \tag{14}$$

In the absence of the hydrodynamic interaction ($\boldsymbol{v} = \boldsymbol{0}$), we obtain $\dot{\phi}_i = F_i/(\zeta R_i) \approx F_i/(\zeta R_{0i}) \equiv \omega_i$, which defines the intrinsic frequency. The radial displacement obeys $R_i(t) - R_{i0} \propto e^{-t/\tau}$, where $\tau = \zeta/k$ is the relaxation time that is typically much shorter than $1/\omega_i$. Therefore, we can assume $R_i = R_{i0}$ and $\dot{\phi} = \omega_i$ in the zeroth order approximation. The flow field is incorporated at the lowest order of the hydrodynamic coupling. On the right hand side of Eq.(11), we can approximate $\boldsymbol{f}_i^{(v)} \approx \zeta\dot{\boldsymbol{r}}_i \approx \zeta R_{i0}\omega_i \boldsymbol{t}_i = F_i \boldsymbol{t}_i$. It gives $\boldsymbol{v}(\boldsymbol{r}_1) \approx \boldsymbol{G}_{12} \cdot \zeta R_{20}\omega_2 \boldsymbol{t}_2 \approx \gamma R_{20}\omega_2 t_{2x} \boldsymbol{e}_x = (\gamma/\zeta) F_2 t_{2x} \boldsymbol{e}_x$, and therefore

$$R_1 = R_{10} + \gamma R_2 \omega_{20} \tau t_{2x} \boldsymbol{e}_x, \tag{15}$$
$$\dot{\phi}_1 = \frac{1}{\zeta R_1} (F_1 + \gamma F_2 t_{2x} t_{1x}) \approx \omega_1 \left[1 + \gamma \left(\frac{R_{20}\omega_2}{R_{10}\omega_1} t_{1x} t_{2x} - \frac{R_{20}\omega_2\tau}{R_{10}} e_{1x} t_{2x}\right)\right] \tag{16}$$

from Eqs.(13,14). The time evolution equation for the phase difference $\Delta = \phi_1 - \phi_2$ is obtained as

$$\dot{\Delta} = \Delta\omega + \gamma \left[\left(\rho\omega_2 - \rho^{-1}\omega_1\right) t_{1x} t_{2x} - \omega_1\omega_2\tau \left(\rho e_{1x} t_{2x} - \rho^{-1} t_{1x} e_{2x}\right)\right], \qquad \Delta\omega = \omega_1 - \omega_2, \quad \rho = \frac{R_{20}}{R_{10}}. \tag{17}$$

Note that the right-hand side of (17) is a function of $\phi_1$ and $\phi_2$ via $t_{ix}$ and $e_{ix}$, the $x$-components of the radial and tangential unit vectors. Assuming $\Delta\omega \ll \omega_i$ and $\gamma \ll 1$, we can average the right-hand side over one period assuming that $\Delta$ is constant during the cycle. This averaging is performed by changing the variables from $\phi_1$, $\phi_2$ to $\Sigma = \phi_1 + \phi_2$ and $\Delta$ and averaging the right-hand side of Eq.(17) over $0 \leq \Sigma < 4\pi$ [6]. This technique allows us to express $\dot{\Delta}$ as a function of $\Delta$ only, which is cast into the form

$$\begin{aligned}
\dot{\Delta} &= \Delta\omega + \gamma \left[ \left(\rho\omega_2 - \rho^{-1}\omega_1\right) \overline{t_{1x}t_{2x}} - \omega_1\omega_2\tau \left(\rho\overline{e_{1x}t_{2x}} - \rho^{-1}\overline{t_{1x}e_{2x}}\right)\right] && (18)\\
&= \Delta\omega + \frac{\gamma}{2} \left[ \left(\rho\omega_2 - \rho^{-1}\omega_1\right) \boldsymbol{q}_1 \cdot \mathcal{R}(\Delta) \cdot \boldsymbol{q}_2 + \omega_1\omega_2\tau \left(\rho + \rho^{-1}\right) \boldsymbol{q}_1 \cdot \mathcal{R}\left(\Delta + \frac{\pi}{2}\right) \cdot \boldsymbol{q}_2 \right] && (19)\\
&= \Delta\omega - C \sin(\Delta - \Delta_0) && (20)\\
&= -\frac{dV}{d\Delta} && (21)
\end{aligned}$$

where $\boldsymbol{q}_i = \boldsymbol{Q}_i \cdot \boldsymbol{e}_x$, $\mathcal{R}(\Delta) = \begin{pmatrix} \cos\Delta & -\sin\Delta \\ \sin\Delta & \cos\Delta \end{pmatrix}$, and the constants $C$ ($> 0$) and $\Delta_0$ are determined by $\gamma$, $\rho$, $\tau$, $\omega_1$, $\omega_2$, $\boldsymbol{q}_1$, and $\boldsymbol{q}_2$. $V(\Delta)$ is an effective potential given by

$$\begin{aligned}
V(\Delta) &= -\Delta\omega \cdot \Delta + \frac{\gamma}{2} \left[ \left(\rho\omega_2 - \rho^{-1}\omega_1\right) \boldsymbol{q}_1 \cdot \mathcal{R}\left(\Delta + \frac{\pi}{2}\right) \cdot \boldsymbol{q}_2 - \omega_1\omega_2\tau \left(\rho + \rho^{-1}\right) \boldsymbol{q}_1 \cdot \mathcal{R}(\Delta) \cdot \boldsymbol{q}_2 \right] && (22)\\
&= -\Delta\omega \cdot \Delta - C \cos(\Delta - \Delta_0). && (23)
\end{aligned}$$

Note that $C$ has the dimension of frequency and is proportional to $h^2/d^3$ via $\gamma$.

### 6 Parameter estimation

Elastic deformations of a bacterial flagellum are characterized by the bending stiffness of the hook and filament. Recent *in vivo* measurements have revealed that the bending rigidity of the hook increases dynamically with the motor torque, reaching values up to $\sim 3.0\,\mathrm{pN} \cdot \mu\mathrm{m}^2$ in the torsionally loaded state [7, 8]. This value is highly comparable to the bending rigidity of the flagellar filament, which is reported to lie in the range of 2–4 $\mathrm{pN} \cdot \mu\mathrm{m}^2$ [9, 10]. Based on these latest findings, we adopted $EI_{\mathrm{hook}} \approx 3.0\,\mathrm{pN} \cdot \mu\mathrm{m}^2$ as a representative value for the hook in the loaded state for our model calculations.

In our bead assay, the 1-$\mu$m beads are attached to the end of truncated flagellar filaments. Considering that the height of the rotational orbit from the surface is $h \sim 1\,\mu\mathrm{m}$ as specified in the *main text*, we assume the truncated filament length to be $L \approx 1\,\mu\mathrm{m}$. The effective spring constant $k$ at the bead position is dominated by the flexibility of the joint and is scaled by the filament length $L$ as:

$$k \approx \frac{EI_{\mathrm{hook}}}{\ell \cdot L^2}, \tag{24}$$

where $\ell \approx 5 \times 10^{-2}\,\mu\mathrm{m}$ is the hook length [11]. Using $EI_{\mathrm{hook}} \approx 3.0\,\mathrm{pN} \cdot \mu\mathrm{m}^2$ and $L \approx 1\,\mu\mathrm{m}$, the effective spring constant is estimated to be $k \approx 60\,\mathrm{pN}/\mu\mathrm{m}$.

The Stokes drag coefficient of the 1-$\mu$m bead in water at 20 °C is $\zeta \approx 1.9 \times 10^{-2}\,\mathrm{pN} \cdot \mathrm{s}/\mu\mathrm{m}$ [12]. Using these values, the relaxation time for the radial displacement is estimated as:

$$\tau = \zeta/k \approx 3 \times 10^{-4}\,\mathrm{s}. \tag{25}$$

This relaxation time ($\sim 0.3\,\mathrm{ms}$) is nearly two orders of magnitude smaller than the typical rotation period ($\sim 20\,\mathrm{ms}$), strongly justifying our theoretical assumption that the bead's radial displacement adjusts rapidly and its position is described by the phase only.

### 7 Validity of the long-distance approximation in experiments

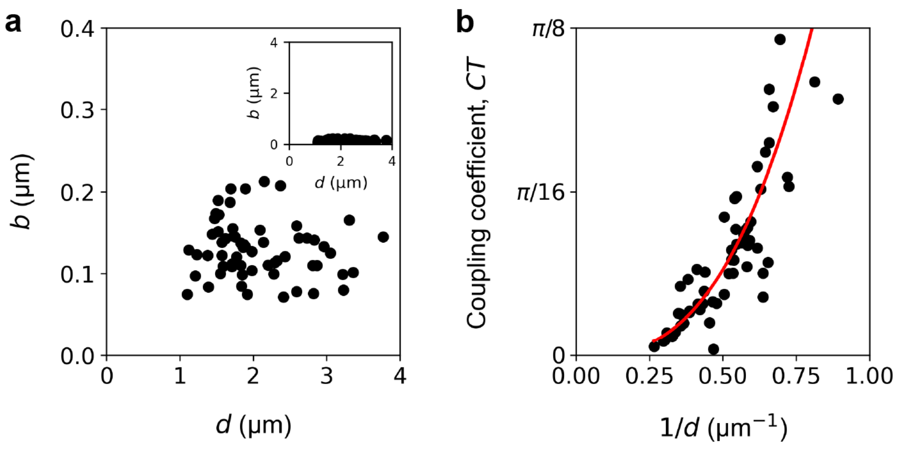


Figure S 6. Verification of the long-distance approximation. Black dots show data from individual bead pairs. (a) Relationship between the inter-bead distance $d$ and the orbital radius $b = (R_{10} + R_{20})/2$ indicates a narrow distribution of $b$ compared to $d$. The inset, plotted with equal vertical and horizontal scales, shows small $b/d$ ratios for all pairs. (b) Dimensionless coupling coefficient $CT$ as a function of $d$. The red line shows a power-law fit, $CT \propto 1/d^n$, which yielded $n = 2.82$ ($R^2 = 0.75$).
